# Establishing Dual-Color Super-Resolution sptPALM for Live-Cell Plasma Membrane Protein Tracking in *Arabidopsis thaliana*

**DOI:** 10.1101/2024.03.13.584811

**Authors:** Leander Rohr, Luiselotte Rausch, Alexandra Ehinger, Nina Glöckner Burmeister, Alfred J. Meixner, Birgit Kemmerling, Klaus Harter, Sven zur Oven-Krockhaus

## Abstract

Super-resolution microscopy techniques have revolutionized cell biology by providing insights into single-molecule dynamics and nanoscale organization within living cells. However, the application of dynamic live-cell methods in plants remains limited by the availability of suitable fluorophores for simultaneous visualization of multiple proteins. To address this challenge, we implemented a dual-color single-particle tracking photoactivated localization microscopy (sptPALM) approach based on codon-optimized photoactivatable fluorescent proteins PA-GFP and PATagRFP. Recently, we demonstrated their individual performance in single-color experiments in *Nicotiana benthamiana* and *Arabidopsis thaliana* cells. Here, we establish their combined use for dual-color sptPALM, enabling the simultaneous tracking of two distinct protein species within the same plant cell. This approach provides a framework to investigate the coordinated dynamics, interactions, and spatial organization of multiple proteins in living plant cells.

## Introduction

Imaging techniques such as super-resolution microscopy provide insights into the dynamics of single molecules and their nanoscale organization of molecular assemblies in vivo, revolutionizing cell biology in recent years [1]. While live-cell single-particle tracking photoactivated localization microscopy (sptPALM) was already performed in the mammalian field by Manley, *et al*. [2] in 2008, the first application in plants was published by Hosy, *et al*. [3] seven years later.

In any biological system, the simultaneous spatiotemporal observation of two differentially labelled proteins is of particular interest to understand the dynamics of proteins within or between membrane nanodomains and their inducible rearrangement at the nanoscale, e.g. oligomeric receptor complexes upon ligand perception. In plants, this type of imaging relies mainly on a limited number of genetically encoded fluorescent proteins (FPs), since the cell wall exacerbates the uptake of external organic dyes commonly used in animal cells [3-6]. For single-color approaches in plants, mEos and its variants proved to be very effective [3, 7-11]. However, mEos-derived FPs are unsuitable for dual-color measurements as their fluorescent native and photoconvertible form block a large part of the spectrum for the excitation and fluorescence readout of other FPs [12]. These experimental hurdles made it previously impossible to simultaneously visualize and track two distinct proteins within the same plant cell.

In our previous work, we screened a range of photoactivatable (PA) FPs based on criteria including spectral compatibility, brightness, maturation, photostability, and blinking behavior. From this analysis, we identified codon-optimized PA-GFP [13] and PATagRFP [14] (Supplementary Table 1) as suitable spectral alternatives to mEos variants and demonstrated that both fluorophores support single-color sptPALM in plant cells without affecting the native dynamics of the fused proteins [7]. Building on these findings, we here demonstrate that PA-GFP and PATagRFP can be used in parallel for dual-color sptPALM, enabling the simultaneous observation of the spatiotemporal behavior of two distinct protein fusions within the same plant cell.

## Results

### Dual-color sptPALM robustly reports RLP44 diffusion in two color channels

To first assess the robustness of sptPALM-based protein diffusion measurements in both fluorescence color channels, the same protein, namely the plasma membrane (PM) located receptor-like protein 44 (RLP44) [15], was used for both PA-GFP and PA-TagRFP fusions. We already demonstrated previously that PA-GFP-, PATagRFP-, and mEos3.2-tagged versions of RLP44 exhibit comparable diffusion coefficients when expressed individually under the control of the native RLP44 promoter in independent transgenic *Arabidopsis thaliana* (*A. thaliana*) lines [7]. The double transgenic line used in this study was obtained by crossing the previously characterized lines expressing RLP44-PA-GFP and RLP44-PATagRFP individually (Supplementary Table 3).

All sptPALM measurements were performed in epidermal hypocotyl cells of seven-day-old, light-grown *A. thaliana* seedlings. The fluorescence signal of the specimen was split into a green and red channel using a dichroic beam splitter, and both channels were projected onto different regions of the same camera sensor via lateral offset to allow separate extraction of the channel data. To exclude tracking artefacts, standard filtering criteria were applied, and data analysis was carried out with OneFlowTraX [7], with datasets comprising 1,500 frames recorded at 20 Hz (for further details, see the Material and Methods section).

As shown in Figure 1a and Figure 1b, both fusion proteins exhibited a comparable spatial distribution within the PM under both mock and osmotic stress conditions (300 mM sorbitol). Osmotic stress treatment was applied as a standard perturbation to modulate membrane protein diffusion and probe the system’s responsiveness to biologically relevant stress, known to alter the dynamics of plant PM proteins [3, 7, 16]. The localization patterns align with previously reported data for RLP44 and other membrane proteins [7, 17].

**Figure 1.**
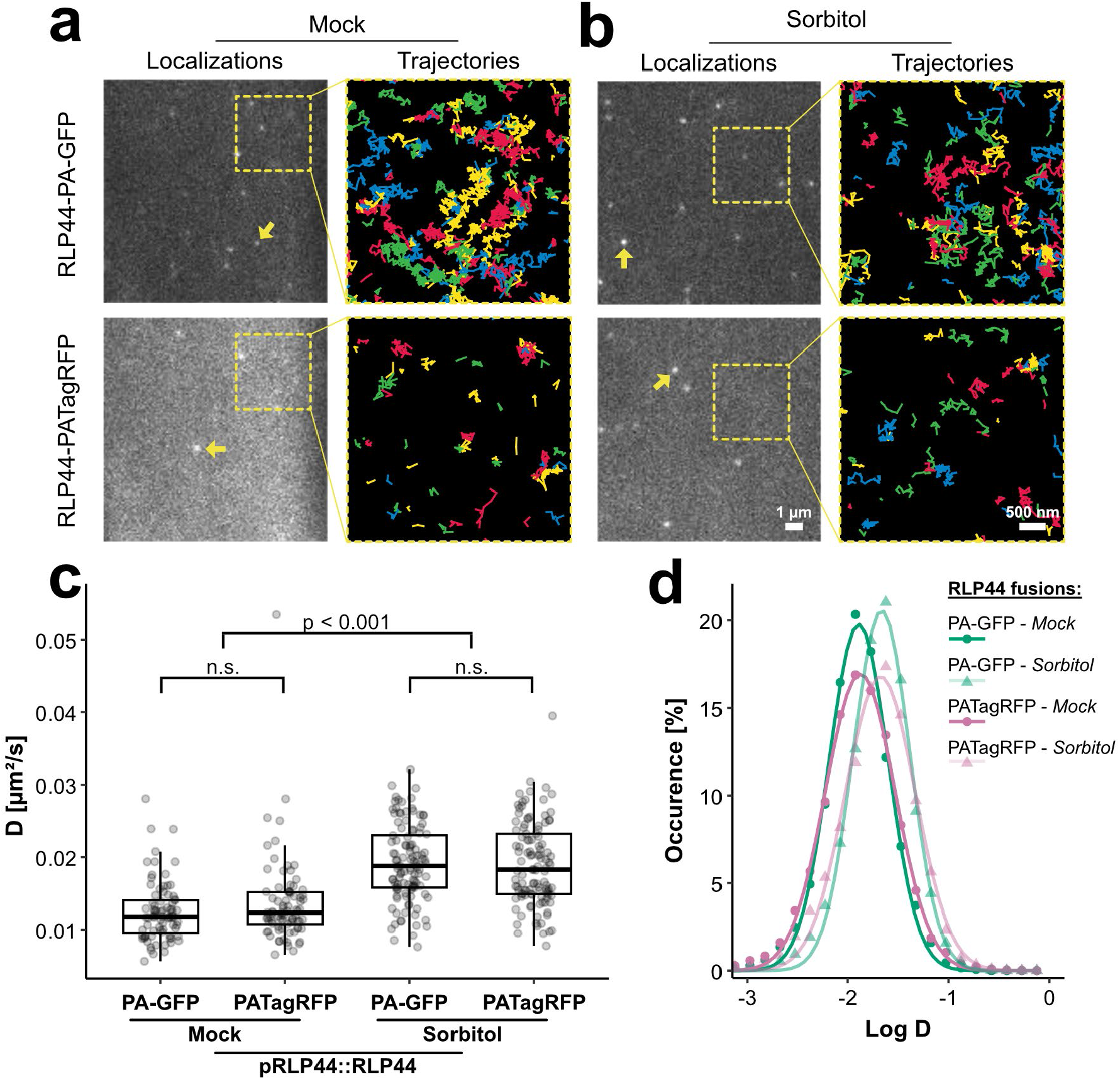
Dual-color sptPALM of RLP44 fused to either PA-GFP or PATagRFP in living *A. thaliana* hypocotyl cells in response to different osmotic conditions. **(a)** Representative sptPALM frames and single-molecule trajectories of RLP44-PA-GFP (top) and RLP44-PATagRFP (bottom) expressed under their native promoter in the plasma membrane of *A. thaliana* hypocotyl cells. Mock-treated cells show diffraction-limited spots (left) and corresponding trajectories (right). Example localizations are indicated by yellow arrows; magnified insets (dashed boxes) highlight areas for the trajectory analysis. **(b)** Analogous depiction of sorbitol-treated cells. **(c)** Peak diffusion coefficients (D) of individual cells derived from Gaussian fits to single-molecule D distributions of each cell for both fusion proteins under mock and 300 mM sorbitol conditions (n ≥ 80 from ≥ 7 plants). Statistics: Shapiro–Wilk test for normality, followed by Kruskal–Wallis Tests. Boxes indicate interquartile range (IQR; 25th and 75th percentiles), the center line represents the median and whiskers extend to 1.5 × IQR. **(d)** log10(D) distributions for RLP44-PA-GFP (green) and RLP44-PATagRFP (magenta) (same raw data as in (c)). Mock conditions are shown as bold curves with points; sorbitol-treated samples as faint curves with triangles.

Absolute diffusion coefficient (D) values were obtained from single-molecule tracks by calculating individual diffusion coefficients and fitting their distribution with a Gaussian function, from which the peak value was extracted, as described previously [3, 7]. RLP44-PA-GFP and RLP44-PATagRFP did not show significant differences under identical osmotic stress conditions (Figure 1c), confirming that the measured diffusion coefficients are independent of the fluorescent tag.

Importantly, we validated the responsiveness of the dual-color system to external perturbations by demonstrating that RLP44 fusions exhibit increased diffusion in both color channels following treatment with 300 mM sorbitol (Figure 1c).

As shown in Figure 1d, the decadic logarithm of D for simultaneously recorded RLP44-PA-GFP and RLP44-PATagRFP fusion proteins displayed a bell-shaped distribution for both osmotic conditions, indicating single RLP44 populations.

These proof-of-principle studies demonstrate that the dual-color system allows simultaneous single-molecule diffusion measurements with identical behavior of both tagged versions in living *A. thaliana* cells, providing a robust and responsive platform for parallel acquisition.

### Extending the dual-color system to two distinct proteins: BRI1 and RLP44

After demonstrating the general applicability of the approach, we next applied dual-color sptPALM to two functionally linked proteins, BRASSINOSTEROID INSENSITIVE 1 (BRI1) and RLP44, which form a ternary complex with BRI1-ASSOCIATED KINASE 1 (BAK1) in the PM [18].

Based on available data we expected lower mobility for BRI1 and higher mobility for RLP44 in our dual-color sptPALM experiments [7]. To test this hypothesis, a double transgenic *A. thaliana* line was generated by crossing lines that independently express BRI1-PA-GFP and RLP44-PATagRFP under the control of their respective native promoters. Both proteins showed a heterogeneous spatial nanoscale distribution within the PM, a characteristic feature of many plant PM proteins [7, 17] (Figure 2a). We observed significantly different diffusion behavior for both BRI1-PA-GFP and RLP44-PATagRFP when expressed within the same cell, with BRI1-PA-GFP displaying significantly lower mobility than RLP44-PATagRFP (Figure 2b).

**Figure 2.**
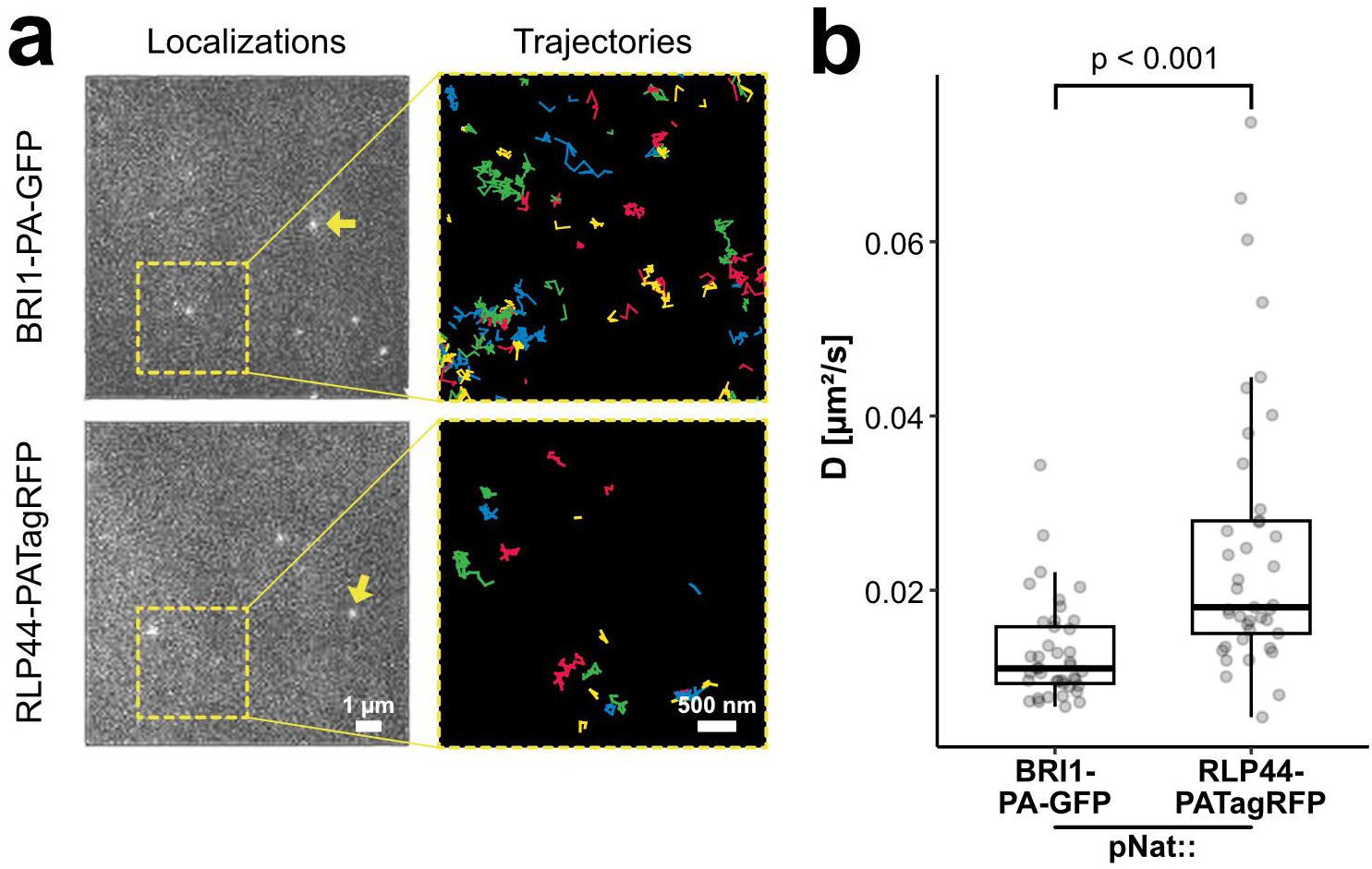
Dual-color sptPALM reveals distinct diffusion behavior of two protein fusions. **(a)** Representative sptPALM frames (left) and single-molecule trajectories (right) of BRI1-PA-GFP (top) and RLP44-PATagRFP (bottom) expressed under their native promoter in the plasma membrane of *A. thaliana* hypocotyl cells. Example localizations are indicated by yellow arrows; magnified insets (dashed boxes) highlight areas for the trajectory analysis. **(b)** Peak diffusion coefficients (D) of individual cells derived from Gaussian fits to single-molecule D distributions of each cell for BRI1-PA-GFP and RLP44-PATagRFP (n = 146 cells from 15 plants). Statistics: Shapiro-Wilk test for normality, followed by Kruskal–Wallis Test. Boxes indicate interquartile range (IQR; 25th and 75th percentiles), the center line represents the median and whiskers extend to 1.5 × IQR.

## Discussion

Approaches based on sptPALM have become increasingly important in recent years, although its application in plant cell biology is still evolving. Early studies and recent advances underscore their significant potential to enhance our understanding of molecular dynamics and spatial organization of proteins at the subcellular level [19].

A key challenge remains multicolor sptPALM, as many signaling processes depend on the coordinated action of at least two proteins [20]. However, the implementation of multicolor approaches has been largely limited by the restricted availability of spectrally compatible and photostable probes, as well as by plant-specific constraints such as the presence of the cell wall and intrinsic autofluorescence [21]. Based on the literature and available data from single-color experiments we selected PA-GFP and PA-TagRFP for the dual-color approach, as they feature well-separated excitation and emission spectra suitable for sptPALM [7].

### Robustness and chromophore independence of dual-color sptPALM

We assessed the robustness of the dual-color approach, including the requirement that dynamic parameters, such as the diffusion coefficient, are independent of the fluorescent probe used. Under identical setup and experimental conditions, the diffusion coefficients derived from PA-GFP– and PA-TagRFP–tagged RLP44 in the same plant cell were in close agreement, demonstrating that neither chromophore-specific photophysical properties nor channel-specific detection biases affect diffusion readouts. An additional important requirement is the analysis of physiological responses to external perturbations, which is critical for comparative studies within a single cellular context. Therefore, we induced osmotic stress with sorbitol, which is known to increase the diffusion coefficient of PM-localized proteins [3, 7, 16]. Consistent with the previous reports, both fluorophore-tagged versions of RLP44 exhibited a comparable increase in diffusion upon osmotic stress in one and the same cell. Together, these proof-of-principle experiments provide strong evidence that dual-color sptPALM can reliably report protein mobility without introducing systematic color-dependent artifacts.

### Discriminating diffusion behaviors of functionally linked proteins

Having validated the technical robustness of the approach, we next applied dual-color sptPALM to two distinct yet functionally connected, spatially associated PM proteins, namely BRI1 and RLP44. The simultaneous observation of both proteins within the same cell allowed a direct comparison of their diffusion behavior under identical experimental and physiological conditions, an advantage that is difficult to achieve with single-color approaches. Consistent with previous reports, BRI1 showed a substantially lower diffusion coefficient than RLP44 in the same PM cells [7]. The ability to resolve these differences within the same environment demonstrates the sensitivity of dual-color sptPALM to intrinsic protein-specific mobility constraints. More importantly, the dual-color approach provides a powerful framework to systematically probe the origins of such mobility differences, for example by applying external cues and directly comparing their effects on associated proteins.

While the qualitative diffusion behaviors observed in our experimental approaches are highly reproducible, slight differences in absolute diffusion coefficients were observed when comparing the values of RLP44 in different experimental contexts. Given the well-documented sensitivity of sptPALM to external parameters such as temperature or the mounting medium [4], these deviations are not unexpected. Importantly, we believe that precisely this sensitivity is an advantage of the dual-color approach: Measuring proteins in the same cellular context minimizes variability and enables more robust studies of protein dynamics and interactions at the nanoscale. Dual-color sptPALM will therefore offer new opportunities for the plant science community to explore complex cellular processes at the nanoscale level.

### Limitations and Outlook: Towards Dynamic Co-Localization Studies

Our dual-color sptPALM approach opens new opportunities to investigate the dynamic organization of signaling complexes at the PM of plant cells. The ability to simultaneously track two proteins within the same cellular context provides the theoretical framework for studies to investigate transient interactions, co-diffusion, or coordinated confinement in response to developmental or environmental cues. At present, however, several methodological limitations constrain the direct extraction of such parameters from dual-color single-molecule data. One major limitation arises from the stochastic nature of PA FPs. Their activation occurs randomly and independently in each channel, making simultaneous detection of two interacting proteins inherently sparse. Consequently, spatial or temporal co-localization events are under-sampled, which can reduce the reliability of downstream analyses such as clustering algorithms that are successfully used in single-color sptPALM plant studies [8, 22, 23]. This challenge is amplified by the opposing demands of sparse localizations for robust tracking versus dense localizations for meaningful clustering analyses. Moreover, additional factors must be considered: in our case, fewer molecules were detected in the red channel, and track lengths differed substantially between differentially tagged PM proteins. While these differences did not affect the calculated diffusion coefficients, reduced detection density and shorter trajectories may limit the robustness of other analyses, such as the identification of nanoscale protein accumulations or the classification of distinct movement states [7, 24, 25]. Thus, caution is warranted when inferring stable molecular interactions solely from spatial proximity or cluster overlap.

It is important to note that these limitations are not specific to plant cells but rather reflect general constraints imposed by the photophysics and stochastic activation of PA fluorophores. Consequently, these limitations apply more widely across cellular systems [26]. Although alternative strategies such as BiFC-PALM [27, 28] have been proposed, these approaches introduce additional limitations that currently hinder their widespread application [27-29].

Looking forward, two major subjects potentially hold promises to overcome these limitations. Firstly, the rapid advancement of artificial intelligence (AI)-based analysis methods in microscopy offers new opportunities to extract meaningful interaction signatures from sparse and complex single-molecule datasets [30, 31]. Machine learning and deep learning approaches are increasingly being applied to classify diffusive motion types and detect transitions between them [32, 33]. Such methods have the potential to yield significant insights, particularly in the context of dual-color analyses aimed at identifying coordinated motion, co-localization, or dynamic clustering. Secondly, ongoing improvements in fluorophore development are expected to further expand the portfolio of suitable candidates for dual-color imaging [34-36]. For applications in the plant cells, especially small, bright, and photostable probes that are capable of efficiently penetrating the plant cell wall are of main interest. Such advances could substantially improve labelling density and synchronization between channels for robust multicolor single-molecule imaging, expanding the scope of dual-color sptPALM toward quantitative co-localization mapping.

In summary, we established dual-color sptPALM as a robust approach for the simultaneous quantification of protein diffusion in living *A. thaliana* cells. Pairing PA-GFP with PATagRFP, we demonstrate that chromophore-dependent bias does not compromise diffusion measurements and that functionally linked but distinct PM proteins can be directly compared within the same cellular context. While current limitations, as discussed above, restrict the direct analysis of clustering behavior or related properties such as molecular interactions, the method provides a solid foundation for future developments. Together with advances in fluorophore design and AI-assisted data analysis, dual-color sptPALM holds strong potential to evolve into a powerful framework for dissecting the dynamic organization and regulation of, for instance, signaling complexes at the plant PM.

## Material and Methods

### Plasmid Construction

All expression clones were constructed using GoldenGate assembly with BB10 as the vector backbone [37]. The promoter sequences were obtained with the help of the Integrated Genome Browser [38]. Level I modules were generated by PCR amplification of the desired sequences and then blunt-end cloned into pJET1.2 (Thermo Fisher Scientific) or ready-to-use obtained by others, such as the pFAST module from Dr. Andrea Gust and the hygromycin resistance module [37]. Fluorophores were designed as C-terminal fusions (D-E module) using a respective linker (Supplementary Table 2). The coding sequence of RLP44 and BRI1 were constructed as B-D modules, eliminating the need for a B-C dummy module. The further procedure was performed as described in Rohr, *et al*. [7]. A full list of used constructs can be found in Supplementary Table 2.

### Plant Material and Growth Conditions

The transgenic stable *A. thaliana* lines expressing one individual fusion protein generated for this study were all in the Columbia (Col-0) background. They were created using the Floral Dip method according to Zhang, *et al*. [39]. The double transgenic plant lines utilized in this study were generated through crossings. Two different lines were studied: (i) Under the control of their native promoter, plants expressing RLP44-PA-GFP-pFAST served as the maternal parent and RLP44-PATagRFP-Hygromycin provided the pollen; (ii) plants transcriptionally controlled by their native promoters, expressing BRI1-PA-GFP-pFAST were used maternally and RLP44-PATagRFP-Hygromycin served as pollen donator.

The resulting seeds were propagated via the presence of the pFAST marker by binocular visual inspection and by selection of survivors on ½ Murashige and Skoog (MS) plates containing 1 % (w/v) sucrose and 0.8 % (w/v) phytoagar supplemented with 25 μM hygromycin. For the presented experiments, lines homozygous for both transgenes were used. For sptPALM measurements, seeds were sterilized with a solution of 70 % ethanol (v/v) and 0.05 % Triton X-100 for 30 minutes followed by a 10-minute treatment with absolute ethanol. Seeds were sown on ½ MS plates (+1 % sucrose and 0.8 % phytoagar) and stratified at 4 °C for at least 24 hours. The light-grown seedlings were cultivated for seven days in growth chambers at 20 °C under long-day conditions (16 hours light / 8 hours dark). A full list of used plant lines can be found in Supplementary Table 3.

### Sample Preparation

For each sptPALM measurement, a seven-day old seedling was placed between two coverslips (Epredia 24×50 mm #1 or equivalent) with a drop of water. This “coverslip sandwich” was then placed on the specimen stage, lightly weighted down by a brass ring. The procedure for the sorbitol (obtained by Roth) treatments was slightly modified. Here, seedlings were incubated in liquid ½ MS medium (+ 1 % sucrose) containing 300 mM sorbitol (or water as mock solution) for five minutes. After this pre-incubation, they were transferred to the coverslip and imaged in the respective incubation solution as mounting medium for up to 20 minutes.

### Microscopy

All sptPALM measurements were performed on a custom-built widefield microscope optimized for variable-angle epifluorescence microscopy (VAEM). The arrangement used for the here described measurements is a modified version of the setup introduced in [7]. In detail: Continuous-wave lasers at 405 nm (iBeam smart 405-S-LP, 100mW, Toptica), 488 nm (Oxxius Laserboxx Diode laser, 100 mW, OXX-488-100, Laser 2000) and 561 nm (Vortran Stradus DPSS laser, 50 mW, VOR-561-050) passed individual clean-up filters (ZET 488/10, F49-488; 560/14 BrightLine HC, F39-561; all AHF analysentechnik AG) and were combined into a single excitation path using beam splitters (Laser Beamsplitter zt 488 RDC, F43-088; Laser Beamsplitter H 405 LPXR, F48-403; all AHF analysentechnik AG), with intensities controlled by an acousto-optic tunable filter (AOTF, Polychromatic Modulator 450 - 700 nm, AOTFnC-VIS-TN, with 4-channel RF driver AA.MPDS4C, both AA OPTO-ELECTRONIC) for rapid switching and precise power adjustment. The lasers were coupled into a single-mode fiber (Polarization-Maintaining FC/PC Fiber Optic Patch Cable PM-S405-XP, P1-405BPM-FC-2, Thorlabs) with an aspheric lens (350 - 700 nm, f = 11.0 mm, NA = 0.25 Aspheric Lens, C220TMD-A, Thorlabs) to ensure a spatially homogeneous beam. The out-coupled beam was collimated (∅1” Achromatic Doublet, ARC: 400-700 nm, f=30 mm, AC254-030-A-ML, Thorlabs) and focused (∅1” UVFS Plano-Convex Lens, f = 150.0 mm, ARC: 350 - 700 nm, LA4874-A-ML, Thorlabs) into the back focal plane of a 100×/1.49 NA oil-immersion objective (Objective alpha Plan-Fluar 100x/1,49 Oil M27, 421190-9800-000, Carl Zeiss Microscopy) after being reflected by a multiband beam splitter (TIRF Quad zt405/488/561/640rpc, F73-410, AHF analysentechnik AG). The VAEM illumination angle was controlled via the lateral translation of a mirror on an electronically controlled linear stage (Right-Angle Prism Dielectric Mirror, MRA20-E02; Linear Stage: 28 mm Travel, ELL17/M; all Thorlabs), included in the excitation pathway after the focusing lens. Low-intensity 405 nm illumination was used for photoconversion or photoactivation when required.

Fluorescence emitted by the sample passed the multiband beam splitter and was focused by a tube lens (1X ZEISS Tube Lens, #13-828, Edmund Optics). After spatial filtering with a rectangular aperture (SP40, Owis), the beam was collimated again (∅1” Achromatic Doublet, ARC: 400-700 nm, f=300 mm, AC254-300-A-ML, Thorlabs) and entered a dual-color split system described in Power et al. (2024). Here, the beam was split into a green and a red channel by a dichroic mirror (Laser-BS H 560 LPXR (T=2mm), F48-558, AHF analysentechnik AG), entering separate arms that were steered and focused individually (∅1” Achromatic Doublet, ARC: 400-700 nm, f=200 mm, AC254-200-A, Thorlabs). They were projected onto the same sCMOS camera (Digital CMOS camera, ORCA-Flash4.0 V2, Hamamatsu Photonics) via a knife-edge mirror (MRAK25-P01, Thorlabs) to introduce a lateral shift between the channel projections. Appropriate emission filters were selected according to the fluorophore (PA-GFP: 488 LP Edge Basic, F76-490; 525/50 BrightLine HC, F37-516; PATagRFP: 561 LP Edge Basic, F76-561; 600/52 BrightLine HC, F39-613; all AHF analysentechnik AG), introduced into the respective arm after the dichroic mirror. In this 4f system, the magnification is adjusted such that one pixel corresponded to 100 nm in the sample plane. Laser power at the sample plane was measured after the objective and adjusted to ensure consistent excitation conditions, and daily dark-frame acquisitions were used for noise correction. Dual-color sptPALM recordings were acquired over 51.2 × 102.4 µm fields of view at 20 Hz, with 1,500 frames collected per movie. The raw data consisted of the green-channel projection on the left half of the camera image and the mirrored red-channel projection on the right half.

### Data Processing and Analysis

A custom-written MATLAB program (The MathWorks Inc., Natick, MA, USA) separated the two channels into individual image stacks and applied mirroring and translational-shift corrections to enable subsequent, independent analysis of each channel.

Data obtained by individual days of measurements were evaluated separately. First, all image stacks originating from the PA-GFP channel were loaded into OneFlowTraX (V1.3) to inspect the quality of the data for each analysis step. The same parameters used previously for one-color experiments were applied [7]. During this process, 128 x 128 pixel regions of interest (masks) were defined within areas containing localizations. The final analysis was then performed in the “Batch analysis” tab of OneFlowTraX. Subsequently, image stacks of the PATagRFP channel were evaluated, using the respective identical masks. Only cells that passed the quality check for both color channels as well as cells that contained n ≥ 10 tracks were included in the final dataset.

Statistical analyses were performed independently for each experimental day using a custom R-Script (Version 4.4.1) to ensure robust behavior. Since different experimental days showed similar behavior, data derived from the same experiment were pooled, their statistics were confirmed, and data were plotted using the ggplot2 [40] and ggpubr [41] packages. The applied statistical test as well as other relevant indices are indicated in the respective figure.

## Author Contributions

LRo: Conceptualization, Data curation, Formal analysis, Investigation, Visualization, Writing – original draft, Writing – review & editing. LRa: Data curation, Formal analysis, Investigation, Writing – review & editing. AE: Resources, Writing – review & editing. NGB: Resources, Writing – review & editing. AJM: Conceptualization, Funding acquisition, Resources, Supervision, Writing – review & editing. BK: Conceptualization, Funding acquisition, Supervision, Writing – review & editing. KH: Conceptualization, Funding acquisition, Supervision, Writing – original draft, Writing – review & editing. SzO-K: Conceptualization, Data curation, Formal analysis, Investigation, Methodology, Software, Writing – original draft, Writing – review & editing.

## Data Availability

The main analysis software OneFlowTraX is freely available for non-commercial use on GitHub: https://github.com/svenzok/OneFlowTraX. Additional custom scripts and code used for data processing and statistical analysis are available from the corresponding author upon reasonable request.

## Acknowledgments and funding

This research was supported by the German Research Foundation (DFG) via the CRC grant 1101 to AJM, BK, KH and SzO-K, and by individual DFG grants to KH (HA 2146/22, HA 2146/23). We also thank the DFG for grants for scientific equipment (FUGG: INST 37/991-1, INST 37/992-1, INST 37/819-1, INST 37/965-1). Furthermore, we would like to thank Dr. Andrea Gust for providing us with the pFAST construct and Jutta Keicher for her help with generating the crossed plant lines.

## Supplementary Information

**Supplementary Table 1.** Nucleotide sequences of the codon-optimized fluorophores PA-GFP and PATagRFP. Codon-optimized nucleotide sequences of the fluorophores PA-GFP and PATagRFP and their respective position. Genes were synthesized by Invitrogen’s GeneArt services (Thermo Fisher Scientific).

| PA-GFP |  |  |  |  |  |  |  |  |  |  |  |  |  |  |  |  |  |
| --- | --- | --- | --- | --- | --- | --- | --- | --- | --- | --- | --- | --- | --- | --- | --- | --- | --- |
| 1 | ATG | GTG | AGC | AAG | GGC | GAA | GAG | TTG | TTC | ACT | GGT | GTT | GTT | CCT | ATC | CTC | GTT |
| 52 | GAG | CTT | GAC | GGT | GAT | GTG | AAC | GGG | CAT | AAG | TTC | TCC | GTT | TCT | GGT | GAA | GGT |
| 103 | GAG | GGT | GAT | GCT | ACT | TAC | GGA | AAG | CTC | ACC | CTC | AAG | TTC | ATC | TGT | ACC | ACT |
| 154 | GGA | AAG | CTC | CCT | GTG | CCT | TGG | CCT | ACT | CTC | GTT | ACC | ACT | TTC | TCT | TAC | GGG |
| 205 | GTG | CAA | TGC | TTC | AGC | AGA | TAC | CCT | GAT | CAT | ATG | AAG | CAG | CAC | GAC | TTC | TTC |
| 256 | AAG | AGC | GCT | ATG | CCT | GAG | GGA | TAC | GTG | CAA | GAG | AGA | ACC | ATT | TTC | TTC | AAG |
| 307 | GAC | GAC | GGG | AAC | TAC | AAG | ACC | AGA | GCT | GAG | GTT | AAG | TTC | GAA | GGT | GAC | ACC |
| 358 | CTC | GTG | AAC | AGG | ATC | GAG | CTT | AAG | GGC | ATC | GAC | TTC | AAA | GAG | GAC | GGA | AAC |
| 358 | ATC | CTC | GGG | CAC | AAG | TTG | GAG | TAC | AAC | TAC | AAC | AGC | CAC | AAC | GTG | TAC | ATC |
| 460 | ATG | GCC | GAC | AAG | CAG | AAG | AAC | GGC | ATC | AAG | GCC | AAC | TTC | AAG | ATC | AGG | CAC |
| 511 | AAC | ATC | GAG | GAT | GGC | TCT | GTT | CAG | CTC | GCT | GAT | CAT | TAC | CAG | CAG | AAC | ACC |
| 562 | CCT | ATT | GGA | GAT | GGA | CCT | GTT | CTT | CTC | CCT | GAC | AAC | CAC | TAC | CTT | AGC | CAC |
| 613 | CAG | AGC | AAG | TTG | AGC | AAG | GAC | CCT | AAT | GAG | AAG | AGG | GAC | CAC | ATG | GTG | CTC |
| 664 | TTG | GAG | TTT | GTT | ACT | GCT | GCT | GGA | ATC | ACC | CTC | GGA | ATG | GAC | GAG | CTT | TAC |
| 715 | AAG | TGA |  |  |  |  |  |  |  |  |  |  |  |  |  |  |  |

| PAtagRFP |  |  |  |  |  |  |  |  |  |  |  |  |  |  |  |  |  |
| --- | --- | --- | --- | --- | --- | --- | --- | --- | --- | --- | --- | --- | --- | --- | --- | --- | --- |
| 1 | ATG | GAA | CTC | ATC | AAA | GAA | AAC | ATG | CAC | ATG | AAG | CTC | TAC | ATG | GAA | GGG | ACC |
| 52 | GTG | AAC | AAC | CAC | CAT | TTC | AAG | TGC | ACA | AGC | GAA | GGT | GAG | GGA | AAG | CCT | TAC |
| 103 | GAG | GGA | ACT | CAG | ACC | ATG | AGA | ATC | AAG | GTG | GTG | GAA | GGT | GGA | CCT | CTT | CCT |
| 154 | TTC | GCC | TTC | GAT | ATT | CTC | GCC | ACC | TCC | TTC | ATG | TAC | GGG | TCC | TCT | ACT | TTC |
| 205 | ATC | AAC | CAC | ACT | CAG | GGA | ATC | CCG | GAC | TTC | TGG | AAG | CAA | TCT | TTT | CCA | GAG |
| 256 | GGA | TTC | ACC | TGG | GAG | AGA | GTG | ACT | ACT | TAC | GAG | GAT | GGT | GGT | GTG | CTC | ACT |
| 307 | GCT | ACT | CAG | GAT | ACT | TCT | CTT | CAG | GAC | GGC | TGC | CTC | ATC | TAC | AAC | GTG | AAG |
| 358 | ATC | AGA | GGT | GTG | AAC | TTC | CCG | TCT | AAC | GGA | CCG | GTG | ATG | AAG | AAA | AAG | ACT |
| 409 | CTC | GGA | TGG | GAG | CCG | TCT | ACC | GAG | AAA | CTT | AAG | CCT | GCT | GAT | GGT | GGA | CTT |
| 460 | GAG | GGA | AGA | GTT | GAC | ATG | GCT | CTT | AAG | CTC | GTT | GGA | GGT | GGA | CAT | CTC | ATC |
| 511 | TGC | AAC | TTC | AAG | ACC | ACC | TAC | AGG | TCT | AAG | AAG | CCG | GCC | AAG | AAC | CTT | AAG |
| 562 | ATG | CCT | GGG | GTT | TAC | TAC | GTG | GAC | AGG | CGT | CTT | GAG | ATT | ATC | AAA | GAG | GCC |
| 613 | GAC | AAA | GAG | ACT | TAC | TGG | GAG | CAG | CAT | GAA | GTG | GCT | GTG | GCT | AGG | TAT | TCT |
| 664 | GAC | CTT | CCA | TCT | AAG | CTC | GGG | CAC | AAG | CTC | AAT | TGA |  |  |  |  |  |

**Supplementary Table 2.**
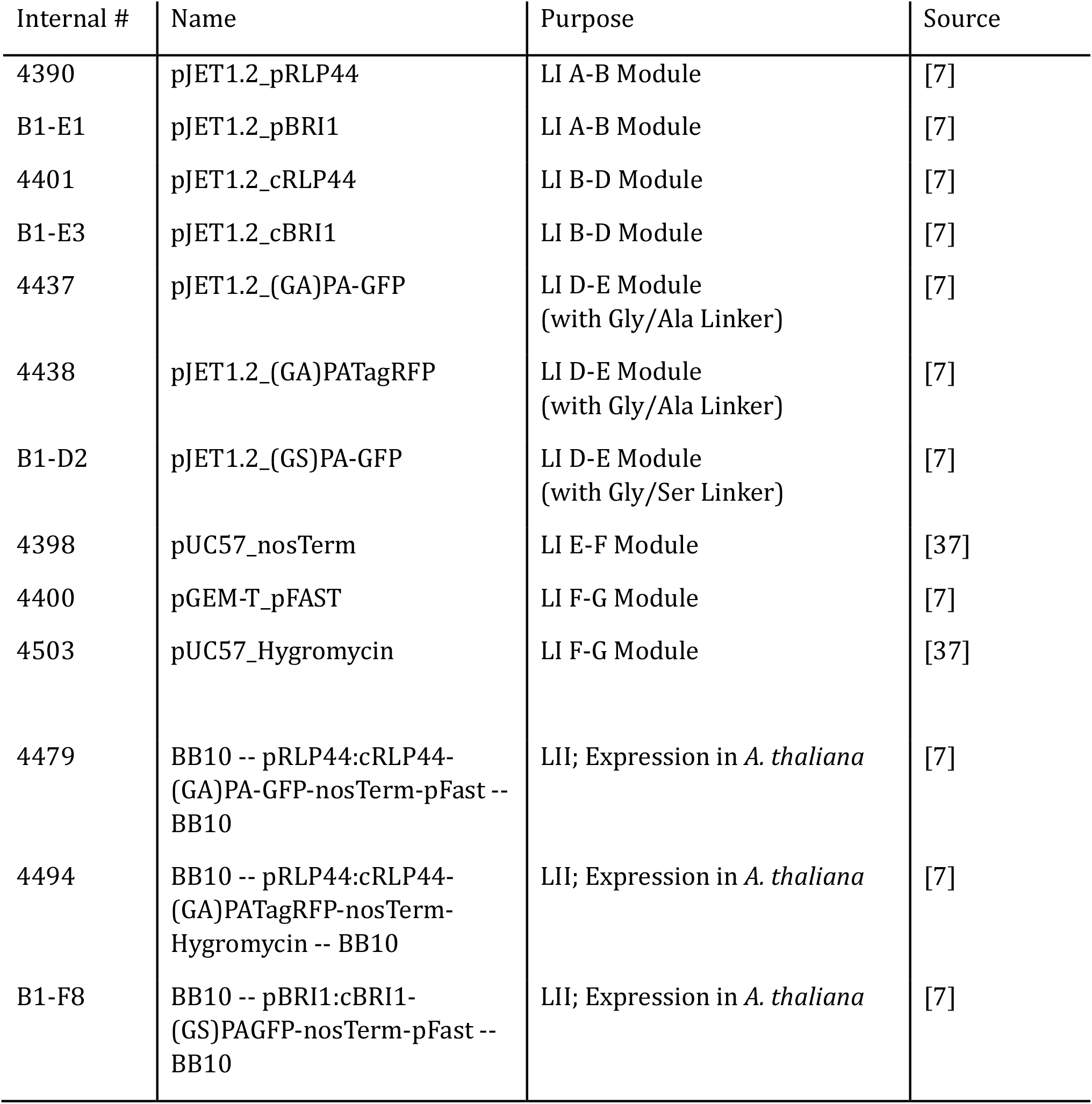
List of constructs. All constructs used are listed with our internal numbering, their name, their purpose and source.

**Supplementary Table 3.**
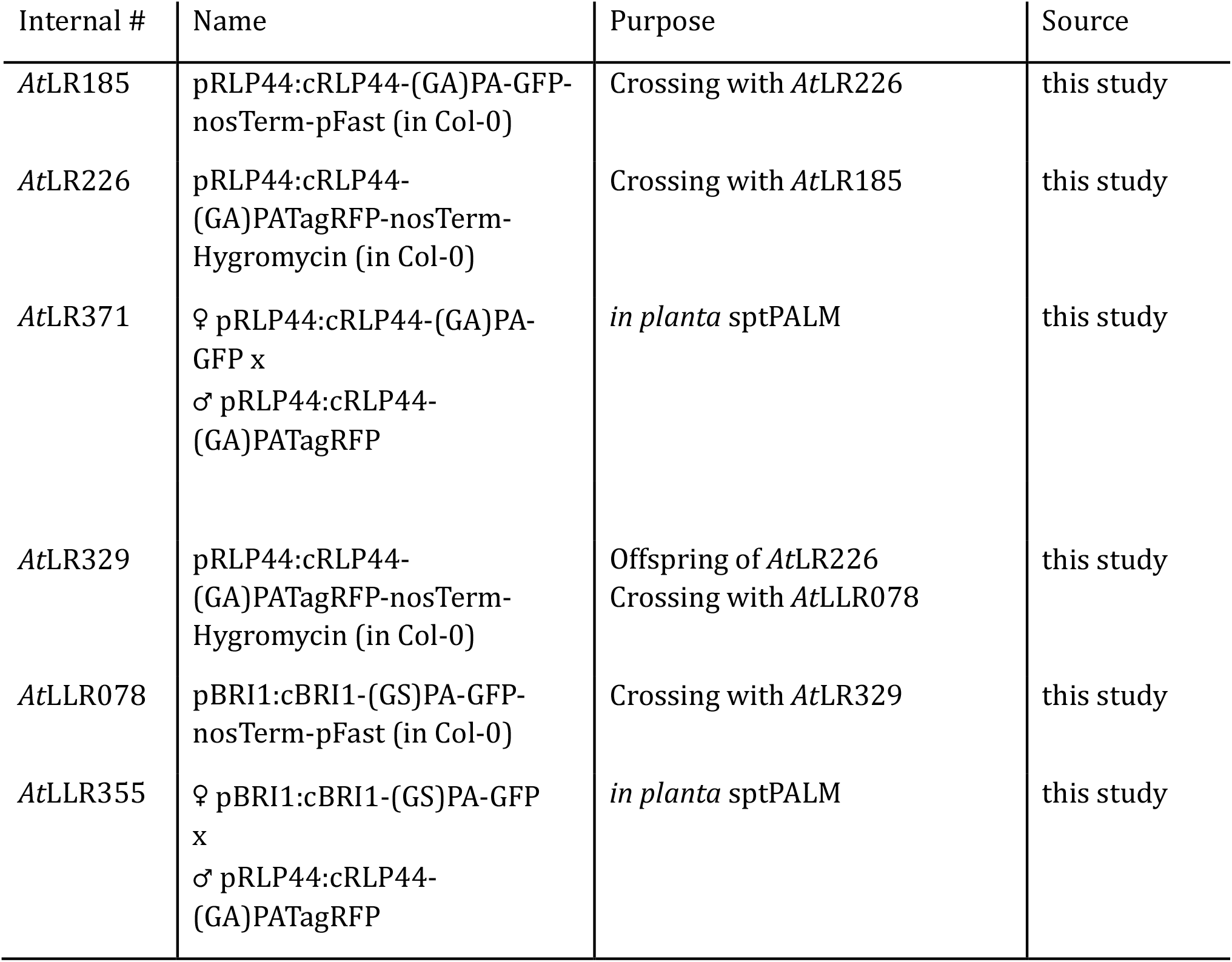
List of used plant lines. All plant lines used are listed with our internal numbering, their name, their purpose and source.

## References

1. Schermelleh, L.; Heintzmann, R.; Leonhardt, H. A guide to super-resolution fluorescence microscopy. J Cell Biol 2010, 190, 165–175 doi: 10.1083/jcb.201002018.

2. Manley, S.; Gillette, J.M.; Patterson, G.H.; Shroff, H.; Hess, H.F.; Betzig, E.; Lippincott-Schwartz, J. High-density mapping of single-molecule trajectories with photoactivated localization microscopy. Nat Methods 2008, 5, 155–157 doi: 10.1038/nmeth.1176.

3. Hosy, E.; Martiniere, A.; Choquet, D.; Maurel, C.; Luu, D.T. Super-resolved and dynamic imaging of membrane proteins in plant cells reveal contrasting kinetic profiles and multiple confinement mechanisms. Mol Plant 2015, 8, 339–342 doi: 10.1016/j.molp.2014.10.006.

4. Bayle, V.; Fiche, J.B.; Burny, C.; Platre, M.P.; Nollmann, M.; Martiniere, A.; Jaillais, Y. Single-particle tracking photoactivated localization microscopy of membrane proteins in living plant tissues. Nat Protoc 2021, 16, 1600–1628 doi: 10.1038/s41596-020-00471-4.

5. Iwatate, R.J.; Yoshinari, A.; Yagi, N.; Grzybowski, M.; Ogasawara, H.; Kamiya, M.; Komatsu, T.; Taki, M.; Yamaguchi, S.; Frommer, W.B.; Nakamura, M. Covalent Self-Labeling of Tagged Proteins with Chemical Fluorescent Dyes in BY-2 Cells and Arabidopsis Seedlings. Plant Cell 2020, 32, 3081–3094 doi: 10.1105/tpc.20.00439.

6. Lelek, M.; Gyparaki, M.T.; Beliu, G.; Schueder, F.; Griffie, J.; Manley, S.; Jungmann, R.; Sauer, M.; Lakadamyali, M.; Zimmer, C. Single-molecule localization microscopy. Nat Rev Methods Primers 2021, 1 doi: 10.1038/s43586-021-00038-x.

7. Rohr, L.; Ehinger, A.; Rausch, L.; Glöckner Burmeister, N.; Meixner, A.J.; Gronnier, J.; Harter, K.; Kemmerling, B.; Zur Oven-Krockhaus, S. OneFlowTraX: a user-friendly software for super-resolution analysis of single-molecule dynamics and nanoscale organization. Front Plant Sci 2024, 15, 1358935 doi: 10.3389/fpls.2024.1358935.

8. Gronnier, J.; Crowet, J.M.; Habenstein, B.; Nasir, M.N.; Bayle, V.; Hosy, E.; Platre, M.P.; Gouguet, P.; Raffaele, S.; Martinez, D.; Grelard, A.; Loquet, A.; Simon-Plas, F.; Gerbeau-Pissot, P.; Der, C.; Bayer, E.M.; Jaillais, Y.; Deleu, M.; Germain, V.; Lins, L.; Mongrand, S. Structural basis for plant plasma membrane protein dynamics and organization into functional nanodomains. Elife 2017, 6 doi: 10.7554/eLife.26404.

9. Mathur, J.; Radhamony, R.; Sinclair, A.M.; Donoso, A.; Dunn, N.; Roach, E.; Radford, D.; Mohaghegh, P.S.; Logan, D.C.; Kokolic, K.; Mathur, N. mEosFP-based green-to-red photoconvertible subcellular probes for plants. Plant Physiol 2010, 154, 1573–1587 doi: 10.1104/pp.110.165431.

10. Platre, M.P.; Bayle, V.; Armengot, L.; Bareille, J.; Marques-Bueno, M.D.M.; Creff, A.; Maneta-Peyret, L.; Fiche, J.B.; Nollmann, M.; Miege, C.; Moreau, P.; Martiniere, A.; Jaillais, Y. Developmental control of plant Rho GTPase nano-organization by the lipid phosphatidylserine. Science 2019, 364, 57–62 doi: 10.1126/science.aav9959.

11. Smokvarska, M.; Bayle, V.; Maneta-Peyret, L.; Fouillen, L.; Poitout, A.; Dongois, A.; Fiche, J.B.; Gronnier, J.; Garcia, J.; Hofte, H.; Nolmann, M.; Zipfel, C.; Maurel, C.; Moreau, P.; Jaillais, Y.; Martiniere, A. The receptor kinase FERONIA regulates phosphatidylserine localization at the cell surface to modulate ROP signaling. Sci Adv 2023, 9, eadd4791 doi: 10.1126/sciadv.add4791.

12. Wiedenmann, J.; Ivanchenko, S.; Oswald, F.; Schmitt, F.; Rocker, C.; Salih, A.; Spindler, K.D.; Nienhaus, G.U. EosFP, a fluorescent marker protein with UV-inducible green-to-red fluorescence conversion. Proc Natl Acad Sci U S A 2004, 101, 15905–15910 doi: 10.1073/pnas.0403668101.

13. Patterson, G.H.; Lippincott-Schwartz, J. A photoactivatable GFP for selective photolabeling of proteins and cells. Science 2002, 297, 1873–1877 doi: 10.1126/science.1074952.

14. Subach, F.V.; Patterson, G.H.; Renz, M.; Lippincott-Schwartz, J.; Verkhusha, V.V. Bright monomeric photoactivatable red fluorescent protein for two-color super-resolution sptPALM of live cells. J Am Chem Soc 2010, 132, 6481–6491 doi: 10.1021/ja100906g.

15. Wolf, S.; van der Does, D.; Ladwig, F.; Sticht, C.; Kolbeck, A.; Schurholz, A.K.; Augustin, S.; Keinath, N.; Rausch, T.; Greiner, S.; Schumacher, K.; Harter, K.; Zipfel, C.; Hofte, H. A receptor-like protein mediates the response to pectin modification by activating brassinosteroid signaling. Proc Natl Acad Sci U S A 2014, 111, 15261–15266 doi: 10.1073/pnas.1322979111.

16. Martiniere, A.; Fiche, J.B.; Smokvarska, M.; Mari, S.; Alcon, C.; Dumont, X.; Hematy, K.; Jaillais, Y.; Nollmann, M.; Maurel, C. Osmotic Stress Activates Two Reactive Oxygen Species Pathways with Distinct Effects on Protein Nanodomains and Diffusion. Plant Physiol 2019, 179, 1581–1593 doi: 10.1104/pp.18.01065.

17. Jaillais, Y.; Ott, T. The Nanoscale Organization of the Plasma Membrane and Its Importance in Signaling: A Proteolipid Perspective. Plant Physiol 2020, 182, 1682–1696 doi: 10.1104/pp.19.01349.

18. Glöckner, N.; Zur Oven-Krockhaus, S.; Rohr, L.; Wackenhut, F.; Burmeister, M.; Wanke, F.; Holzwart, E.; Meixner, A.J.; Wolf, S.; Harter, K. Three-Fluorophore FRET Enables the Analysis of Ternary Protein Association in Living Plant Cells. Plants (Basel) 2022, 11doi: 10.3390/plants11192630.

19. Zur Oven-Krockhaus, S.; Rohr, L.; Rausch, L.; Harter, K. Revealing plasma membrane protein dynamics in living plant cells with single-molecule tracking. J Exp Bot 2025, 77, 178–188 doi: 10.1093/jxb/eraf417.

20. McWhite, C.D.; Papoulas, O.; Drew, K.; Cox, R.M.; June, V.; Dong, O.X.; Kwon, T.; Wan, C.; Salmi, M.L.; Roux, S.J.; Browning, K.S.; Chen, Z.J.; Ronald, P.C.; Marcotte, E.M. A Pan-plant Protein Complex Map Reveals Deep Conservation and Novel Assemblies. Cell 2020, 181, 460–474 e414 doi: 10.1016/j.cell.2020.02.049.

21. Donaldson, L. Autofluorescence in Plants. Molecules 2020, 25 doi: 10.3390/molecules25102393.

22. Danek, M.; Hdedeh, O.; Amo, J.; Boutet, J.; Neubergerova, M.; Safi, H.; Abuzeineh, A.; Martin-Barranco, A.; Fiche, J.B.; Mercier, C.; Dumortier, B.; Krouk, G.; Nollmann, M.; Pleskot, R.; Boutte, Y.; Santoni, V.; Mongrand, S.; Martiniere, A.; Zelazny, E. Mechanisms controlling the plasma membrane targeting and the nanodomain organization of the plant SPFH protein HIR2. Plant J 2026, 126, e70879 doi: 10.1111/tpj.70879.

23. Weber, H.; Ehinger, A.; Kolb, D.; Fallahzadeh-Mamaghani, V.; Halter, T.; Franz-Wachtel, M.; Zur Oven-Krockhaus, S.; Gronnier, J.; Zipfel, C.; Harter, K.; Kemmerling, B. Arabidopsis HYPERSENSITIVE INDUCED REACTION 2 affects plasma membrane 1 receptor pathways and organization. bioRxiv 2025, doi: 10.1101/2025.04.11.648320.

24. Rohr, L.; Rausch, L.; Harter, K.; Zur Oven-Krockhaus, S. Contrasting Effects of Cytoskeleton Disruption on Plasma Membrane Receptor Dynamics: Insights from Single-Molecule Analyses. bioRxiv 2024, doi: 10.1101/2024.09.09.612020.

25. von Arx, M.; Xhelilaj, K.; Schulz, P.; Zur Oven-Krockhaus, S.; Gronnier, J. Photochromic reversion enables long-term single-molecule tracking in living plants. bioRxiv 2026, doi: 10.1101/2024.04.10.585335.

26. Vojnovic, I.; Winkelmeier, J.; Endesfelder, U. Visualizing the inner life of microbes: practices of multi-color single-molecule localization microscopy in microbiology. Biochem Soc Trans 2019, 47, 1041–1065 doi: 10.1042/BST20180399.

27. Liu, Z.; Xing, D.; Su, Q.P.; Zhu, Y.; Zhang, J.; Kong, X.; Xue, B.; Wang, S.; Sun, H.; Tao, Y.; Sun, Y. Super-resolution imaging and tracking of protein-protein interactions in sub-diffraction cellular space. Nat Commun 2014, 5, 4443 doi: 10.1038/ncomms5443.

28. Nickerson, A.; Huang, T.; Lin, L.J.; Nan, X. Photoactivated localization microscopy with bimolecular fluorescence complementation (BiFC-PALM) for nanoscale imaging of protein-protein interactions in cells. PLoS One 2014, 9, e100589 doi: 10.1371/journal.pone.0100589.

29. Ren, H.; Ou, Q.; Pu, Q.; Lou, Y.; Yang, X.; Han, Y.; Liu, S. Comprehensive Review on Bimolecular Fluorescence Complementation and Its Application in Deciphering Protein-Protein Interactions in Cell Signaling Pathways. Biomolecules 2024, 14 doi: 10.3390/biom14070859.

30. Khater, I.M.; Nabi, I.R.; Hamarneh, G. A Review of Super-Resolution Single-Molecule Localization Microscopy Cluster Analysis and Quantification Methods. Patterns (N Y) 2020, 1, 100038 doi: 10.1016/j.patter.2020.100038.

31. Munoz-Gil, G.; Bachimanchi, H.; Pineda, J.; Midtvedt, B.; Fernandez-Fernandez, G.; Requena, B.; Ahsini, Y.; Asghar, S.; Bae, J.; Barrantes, F.J.; Bender, S.W.B.; Cabriel, C.; Conejero, J.A.; Escoto, M.; Feng, X.; Haidari, R.; Hatzakis, N.S.; Huang, Z.; Izeddin, I.; Jeong, H.; Jiang, Y.; Kaestel-Hansen, J.; Mine-Hattab, J.; Ni, R.; Park, J.; Qu, X.; Saavedra, L.A.; Sha, H.; Sokolovska, N.; Zhang, Y.; Volpe, G.; Lewenstein, M.; Metzler, R.; Krapf, D.; Volpe, G.; Manzo, C. Quantitative evaluation of methods to analyze motion changes in single-particle experiments. Nat Commun 2025, 16, 6749 doi: 10.1038/s41467-025-61949-x

32. Chen, Z.; Geffroy, L.; Biteen, J.S. NOBIAS: Analyzing anomalous diffusion in single-molecule tracks with nonparametric Bayesian inference. Front Bioinform 2021, 1 doi: 10.3389/?binf.2021.742073.

33. Kowalek, P.; Loch-Olszewska, H.; Szwabinski, J. Classification of diffusion modes in single-particle tracking data: Feature-based versus deep-learning approach. Phys Rev E 2019, 100, 032410 doi: 10.1103/PhysRevE.100.032410.

34. Gaytan, P.; Roldan-Salgado, A. Photoactivatable Blue Fluorescent Protein. ACS Omega 490 2024, 9, 28577–28582 doi: 10.1021/acsomega.4c02603.

35. Pak, Y.L.; Jing, Y.; Wu, W.; Guo, Y.; Wang, Z.; Liu, G.; Song, J. Recent Advances in Single-Molecule Localization Based Super-Resolution Imaging. Small 2026, 22, e10238 doi: 10.1002/smll.202510238.

36. Shcherbakova, D.M.; Sengupta, P.; Lippincott-Schwartz, J.; Verkhusha, V.V. Photocontrollable fluorescent proteins for superresolution imaging. Annu Rev Biophys 2014, 43, 303–329 doi: 10.1146/annurev-biophys-051013-022836.

37. Binder, A.; Lambert, J.; Morbitzer, R.; Popp, C.; Ott, T.; Lahaye, T.; Parniske, M. A modular plasmid assembly kit for multigene expression, gene silencing and silencing rescue in plants. PLoS One 2014, 9, e88218 doi: 10.1371/journal.pone.0088218.

38. Freese, N.H.; Norris, D.C.; Loraine, A.E. Integrated genome browser: visual analytics platform for genomics. Bioinformatics 2016, 32, 2089–2095 doi: 10.1093/bioinformatics/btw069.

39. Zhang, X.; Henriques, R.; Lin, S.S.; Niu, Q.W.; Chua, N.H. Agrobacterium-mediated transformation of Arabidopsis thaliana using the floral dip method. Nat Protoc 2006, 1, 641–646 doi: 10.1038/nprot.2006.97.

40. Wickham, H. ggplot2. Use R! 2016, doi:10.1007/978-3-319-24277-4.

41. Kassambara, A. ggpubr: ggplot2 Based Publication Ready Plots. Available online: 10.32614/CRAN.package.ggpubr (accessed on 2026-05-26).

